# A 3D printed mini-gel electrophoresis system for rapid and inexpensive DNA nanoswitch biosensing

**DOI:** 10.64898/2026.01.21.700818

**Authors:** Vinod Morya, Andrew Hayden, Lifeng Zhou, Dadrian Cole, Ken Halvorsen

## Abstract

Gel electrophoresis has been a cornerstone laboratory technique for decades, yet it is often viewed as cumbersome, costly, and has remained confined to laboratory settings. Recent advances in DNA nanotechnology have repurposed electrophoresis as a primary readout for some biosensing applications such as DNA nanoswitches, where a conformational change in a DNA structure indicates the presence of a target molecule. Conventional gel electrophoresis setups not ideal for such targeted applications, with moderate equipment cost, excessive reagent use, and time-consuming processes. Here, we adopt a reductionist, application-driven approach to redesign gel electrophoresis specifically for DNA nanoswitch-based detection. We present a fully 3D-printable mini gel electrophoresis system that incorporates conductive plastic electrodes, demonstrating performance comparable to conventional systems using platinum electrodes. By optimizing the inter-electrode distance and running parameters, our system resolves the on/off states of DNA nanoswitches in as little as one minute. We further show that the device operates reliably at low voltages, including when powered by a USB power bank, and even enables instrument-free nanoswitch readout using an LED with a cell-phone camera. Our design substantially reduces the cost, voltage requirements, material usage, operational complexity, and experiment time. These improvements make gel-based biosensing more practical outside traditional laboratory environments, paving the way for broader adoption of gel electrophoresis in point-of-care and resource-limited settings.

## Introduction

Gel electrophoresis is a common and foundational laboratory technique for separating charged polymers by size, topology, or conformation. Perhaps the most widely used format is agarose gel electrophoresis of DNA and RNA, which has remained largely unchanged since its introduction in 1972.^1^ The method is deeply integrated into molecular biology workflows, including cloning, PCR analysis, genetic identification, and nucleic acid quality assessment.^2,3^ Despite its ubiquity, the typical equipment has seen minimal evolution. Standard laboratory gel boxes rely on large hand-cast slab gels (often ∼50–200 cm^2^), paired with sizable buffer reservoirs (∼0.4–2 L), and require relatively high voltages (50–150 V) delivered by benchtop power supplies^4^. Visualization of the separated nucleic acids typically relies on intercalating dyes and costly gel documentation systems, which adds expense and operational constraints. Together, these elements make traditional gel electrophoresis confined to laboratory settings, moderately expensive to run, and a source of substantial chemical, and electrical waste. Remarkably, commercial gel boxes are still strikingly similar to designs introduced more than half a century ago^5,6^.

Still, electrophoresis has experienced some innovation, including advancements in materials, miniaturization, microfluidic electrophoresis, and capillary electrophoresis systems.^7–10^ However, these innovations have largely emerged to address analytical performance at the research or clinical scale, rather than to rethink the basic accessibility, size, or operability of the standard slab-gel format. Little effort has focused on redesigning agarose electrophoresis for simplicity, portability, or low-resource settings. Given how entrenched gel electrophoresis is in biological workflows, the lack of optimization for field use, rapid diagnostics, or consumer-level operation is notable.

In recent years, electrophoresis has also taken on a new role as a primary readout for some biosensing assays.^11–13^ Rather than merely separating nucleic acids, electrophoresis can directly report on molecular detection events through mobility shifts. This shift in usage opens the possibility that electrophoresis could support low-cost diagnostic platforms, if the technique itself can be simplified and adapted for point-of-care (POC) contexts.

We have previously developed DNA nanoswitches that use gel electrophoresis as a readout for detecting a wide range of biomolecules.^14–18^ DNA nanoswitches operate by converting a molecular recognition event into a topological change in the DNA structure (Figure 1a).^19^ When the nanoswitch binds its target, it undergoes a conformational change, shifting from a linear to a looped state, producing a clear mobility shift during electrophoresis (Figure 1b). Importantly, the platform is fully programmable and has been adapted to detect diverse classes of biomarkers, including DNA^19^, various RNA species^14,16^, proteins^20^, and combinations of molecular targets^21^. The nanoswitch system is highly cost-effective (typically <1 cent per nanoswitch), versatile, and sensitive. Despite these advantages, reliance on traditional gel electrophoresis has been one of the main criticisms of nanoswitch-based detection, particularly for use in point-of-care diagnostics or resource-limited environments.

**Figure 1:**
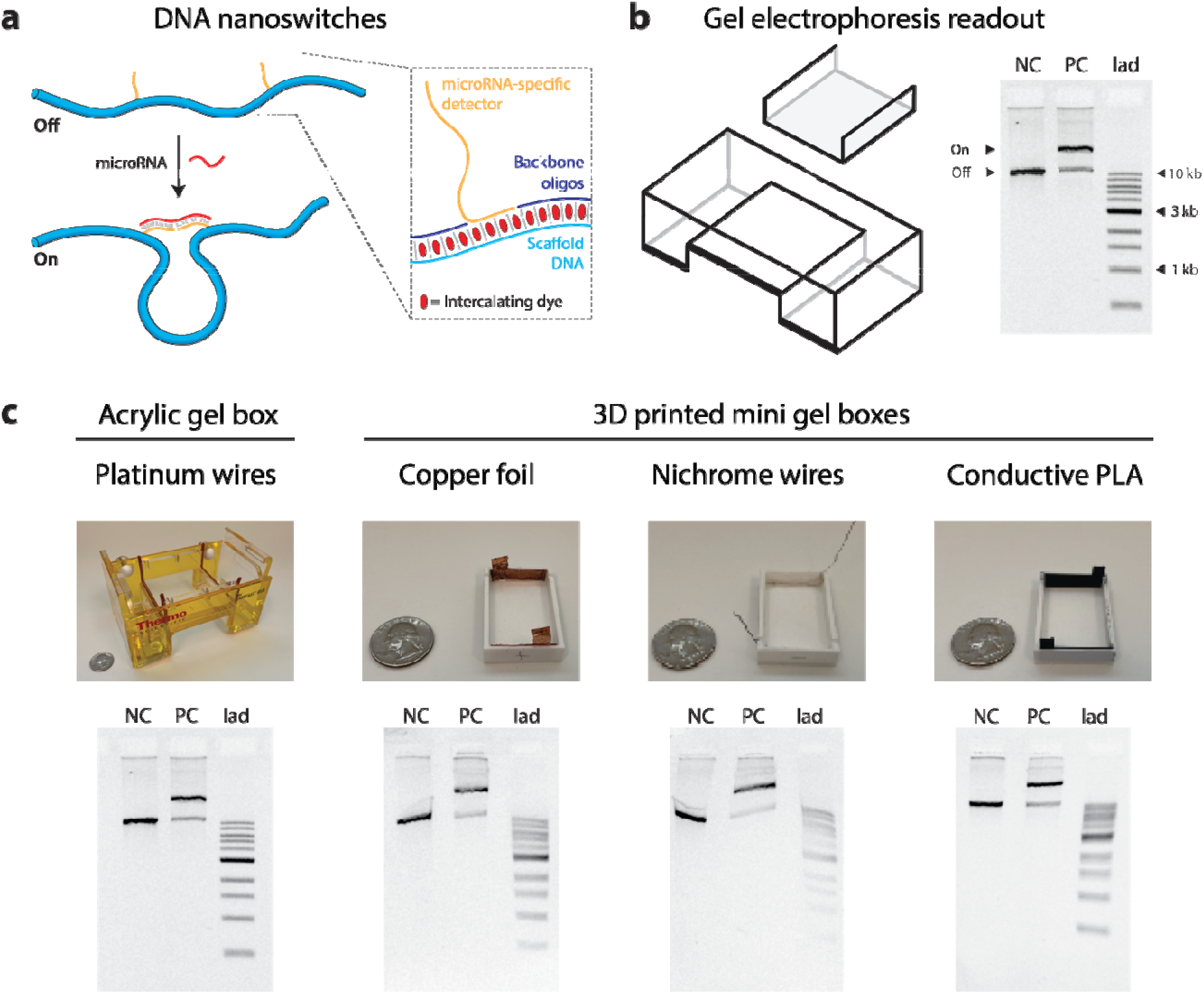
DNA nanoswitch concept and development of custom gel boxes. a) DNA nanoswitches are designed to reconfigure from linear to looped in the presence of a molecular target. B) Agarose gel electrophoresis enables identification of the linear (unlooped) and looped forms. C) Comparison of a standard gel setup with custom made 3D printed mini-gel boxes using copper, Nichrome, or conductive PLA electrodes. All gels were run at 5.5 V/cm for 45 minutes. (US quarter for scale)

Here, we set out to streamline and re-engineer gel electrophoresis as a dedicated readout for DNA nanoswitches. The mobility shift between the on and off states is large enough that full-scale electrophoresis is unnecessary. We first miniaturized the gel system to allow lower-voltage operation, decrease running time, and reduce cost of both manufacturing and operation. We evaluated various electrode materials and found the best performance and cost effectiveness using a fully 3D-printed design with conductive plastic electrodes. With this system, we demonstrate nanoswitch detection in just a few minutes at voltages compatible with standard battery or USB power bank operation. Finally, we evaluated dyes, lighting strategies, and low-cost optical filters to enable imaging with a cell-phone camera or even visible detection by eye without the need for specialized instruments. We anticipate that these efforts represent an early step toward a new generation of gel-based point-of-care assays for healthcare, environmental monitoring, and low-resource diagnostics.

## Results

Our primary goal in this work was to reimagine gel electrophoresis as a readout for DNA nanoswitches – particularly to optimize the system to reduce cost, time, voltage, and experimental effort. We took a reductionist approach to determine the minimal requirements, and the first consideration was that resolving our DNA nanoswitches requires only a small fraction of the area of a typical gel. The smallest commercial mini gel system we use (Owl B1A) is still overkill for our application, with gel size of 7 x 8 cm, a footprint of about 15 x 10 cm, and holding ∼400 mL of running buffer. In contrast our DNA nanoswitch bands occupy only ∼ 0.5 x 0.5 cm (less than 1% of the area) for a single lane. We considered that a smaller gel footprint could offer many potential advantages including closer electrodes for lower voltage operation, less reagent use including agarose and running buffer, and lower overall equipment cost. To evaluate a smaller gel design, we 3D printed a plastic (PLA) tray and comb to accommodate 3 gel lanes in a 40 mm long gel box (**Figure 1c**). Based on our previous use of buffer less gel electrophoresis (E-gel system, Thermo Fisher), we hypothesized that a large buffer reservoir was unnecessary for our application and that a few mL of buffer to overlay the gel would suffice.

Most standard gel electrophoresis systems use platinum wires, which are known for high conductivity and low corrosion but also extremely high cost. For our mini gel box, we considered three lower cost materials to use as electrodes – copper foil, nichrome wire, and conductive plastic (**Figure 1c**). We found all three materials worked, and we were pleasantly surprised to see that conductive PLA had the straightest bands and best resolution among the three. The conductive PLA is also the easiest to work with since it can be entirely 3D printed and did not exhibit visible surface fouling due to oxidation as the copper and nickel wires did. Overall, we found the performance of our 3D printed mini-gel with conductive plastic electrodes and no buffer reservoirs performed similarly to our conventional gel system with platinum electrodes that we used as our gold standard benchmark.

Considering the good performance and versatility of our 3D mini-gel design, we decided to move forward with this design for further optimization. First, we tested whether a buffer gap at each electrode was beneficial, by running identical gels with or without the gap (**Figure S1**). We found that both systems produced suitable results, but we noticed that a small gap helped reduce entrapment of bubbles generated at the electrodes, potentially helping to reduce the pH stress and allow better heat dissipation. Based on this we decided to include a 1 mm gap between the gel and each electrode for future studies. Next, we assessed repeatability by printing 5 identical gel boxes and running fresh gels in each one (**Figure S2**). We also assessed re-use of the same gel box multiple times, by running 5 sequential gels in one box. We found that the performance degraded slightly over time but still produced acceptable results after 5 runs. We also compared the effectiveness of 3D printed combs with acrylic comb (**Figure S3**). We found that the 3D printed combs produce indistinguishable results from the acrylic comb, and all print qualities also gave similar results.

A primary advantage of 3D printing is rapid and inexpensive prototyping. We took advantage of this capability to evaluate the inter-electrode distance of our 3D printed mini-gel, which is not a parameter that can be changed easily in standard lab electrophoresis. Since the driving force for electrophoresis comes from the electric field strength (i.e. nominally the voltage divided by the inter-electrode distance), moving the electrodes closer should allow for separation at lower total voltages. We designed and 3D printed gel boxes with inter-electrode distances ranging from 10 mm to 40 mm, and ran gels keeping a constant nominal electric field (**Figure 2**). Migration in the largest boxes of 30-40 mm was very similar to the Owl reference gel, while the smaller gel boxes started to exhibit some band compression (**Figure 2b**). In principle a constant electric field would produce uniform migration across all gel boxes, so these results suggest some confounding factors that are likely changing field strength. For our DNA nanoswitch studies, we identified the 15 and 20 mm spacings to be the best mix of low voltage and a box and gel still large enough to physically handle without much trouble.

**Figure 2:**
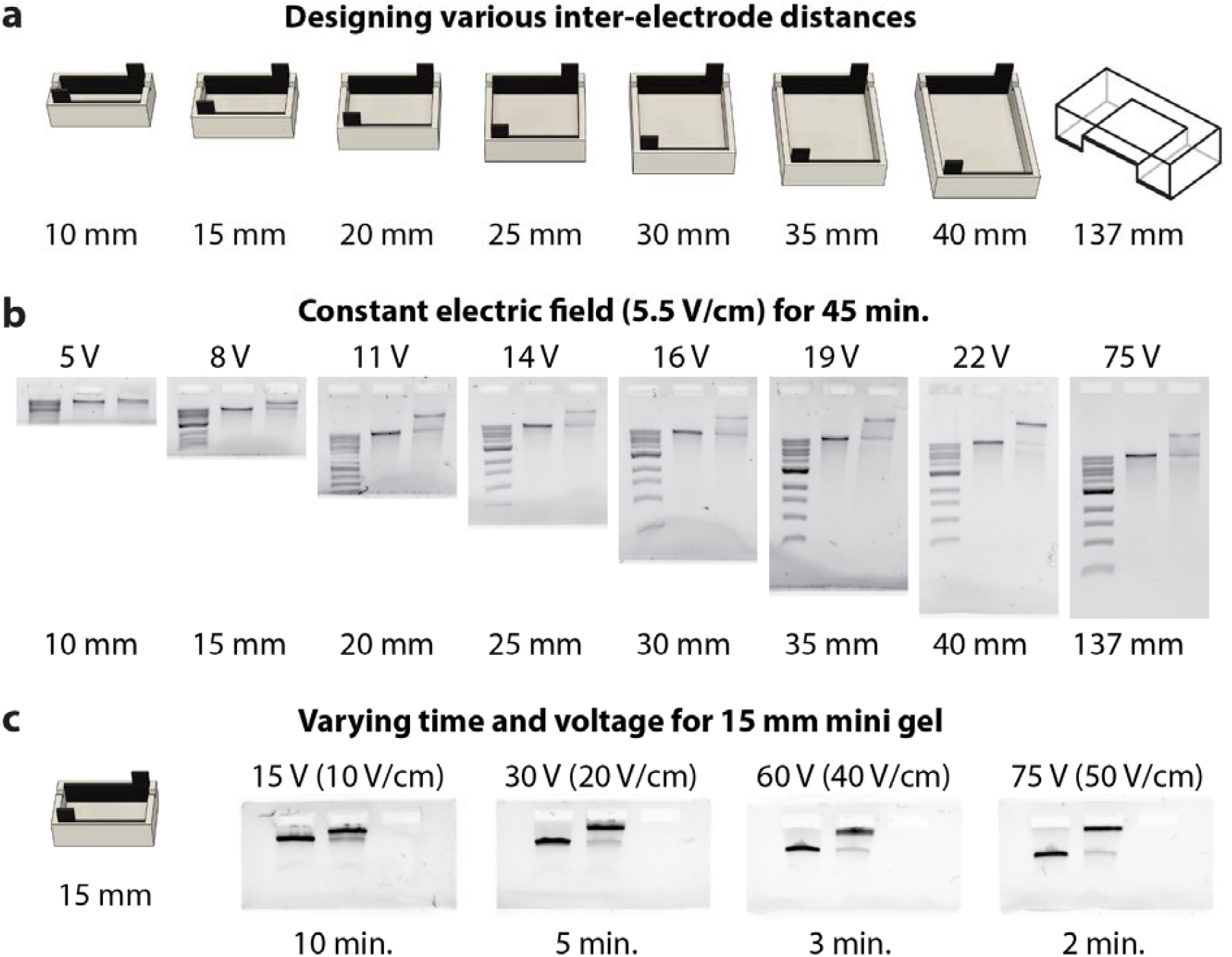
Optimization of electrode spacing. a) Cartoon showing how reducing interelectrode length and voltage can maintain the same electric field and DNA mobility. b) Design and testing of mini-gel boxes with various interelectrode distances and comparisons against a commercially available gel box. Lanes in all the gels from left to right: 1= 1kb DNA ladder, 2= negative control, 3=positive control. c) Testing different time and voltages for 15 mm minigel box. Lanes in all the gels from left to right: 1= negative control, 2=positive control, 3= empty.

Proceeding with the 15 mm design, we further optimized the running conditions with the goal of achieving separation in just a few minutes. We first tested different gel percentages with constant running conditions and found relatively similar results across percentages between 0.7 and 1.2%. We chose 0.8% for further experiments, as it gives better balance between band resolution and agarose used (**Figure S4**). We next systematically increased the applied voltage while proportionally decreasing the run time to evaluate the minimum time required to achieve band resolution. (**Figure 2c**). We found that we could run the gel with electric field strengths as high as 50 V/cm, allowing the DNA nanoswitches to be resolved in as little as 2 minutes. It is interesting to note that this field strength would correspond to >650V in our Owl mini box, outside of the recommended operating range of the Owl box and of many voltage supplies.

Moving toward use outside of a lab environment, we considered which voltages would be easily accessible by battery power for non-lab use. We tested a 1.5 V AAA battery (4 in series, 6V), 1.5 V coin cell (4 in series, 6V), and a 9V battery. We were able to resolve our nanoswitches using all of these power sources, with varying time requirements (**Figure 3a,b**). The use of portable batteries eliminates the requirement for costly standard benchtop power supply systems. To further remove the need for expensive and bulky gel imaging systems, we demonstrate that SYBR Safe staining, in combination with a blue-light source and an orange safety filter, enables gel detection using a standard cell phone camera (**Figure 3c,d**). To bring all of these innovations together, we demonstrate end-to-end detection of a rRNA from total RNA extracted from HeLa cells (**Figure 3e**). We tailored our nanoswitch to detect 5S rRNA, by replacing the detector oligos at V4 and V8 positions. A synthetic DNA mimic or 5S rRNA was used as positive control and the total RNA sample (250 ng/µl) isolated from HeLa cells was used as a test sample.

**Figure 3:**
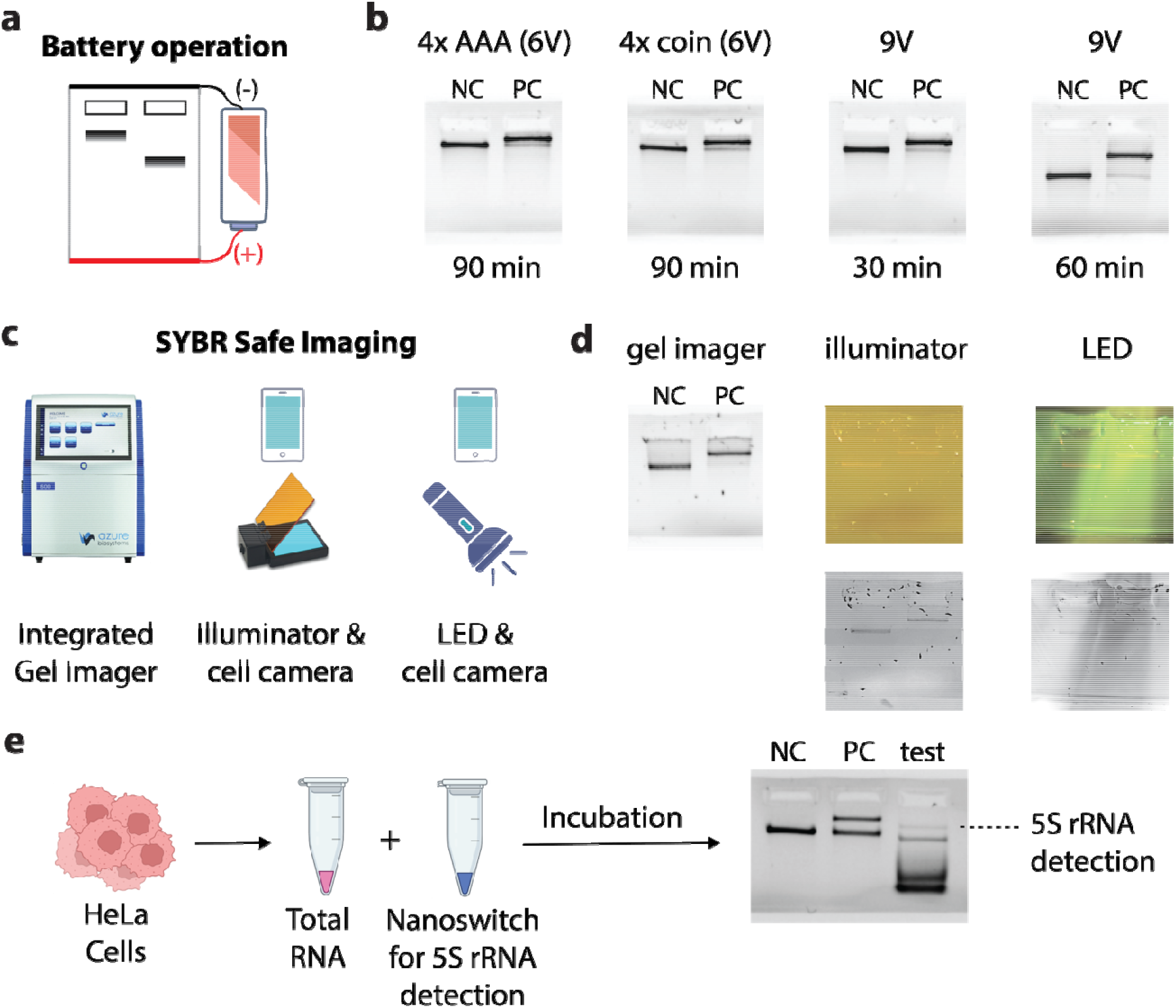
Instrument-free operation and imaging; and analyzing a total RNA sample. a) Mini gels allow low voltage operation compatible with battery power, b) Three different power sources were tested, with gel results at various times, d) cartoon of required components for gel imaging, d) A single gel imaged with three variants of gel imaging, e) Detection of 5S rRNA in total RNA sample isolated from HeLa cells in a 15mm Minigel box, Lane 1= NC (negative control), Lane 2= PC (positive control), Lane 3= test (total RNA sample).

## Discussion

Here we have shown that gel electrophoresis systems can be re-designed and rapidly and inexpensively manufactured to suit the needs of specific applications such as biosensing. For our DNA nanoswitches, we demonstrate that fully 3D printed mini-gel boxes can save money, time, and materials for identifying biomolecules when compared with commercially available gel systems. For the nanoswitch assay in particular, we have shown that we can get the same result with ∼1000x lower equipment cost, ∼100x lower buffer volume, ∼10x lower voltage, ∼10x lower reagent cost, and ∼10x lower time. Our solution is transformative enough to potentially turn thousands of dollars of equipment into low-cost consumables.

Agarose gel electrophoresis as a laboratory technique has been relatively unchanged in nearly 50 years. Commonly used gel boxes available today from lab equipment suppliers are practically indistinguishable from the first one presented in 1977^6^. There have been some progression of techniques such as the advent of capillary gel electrophoresis^22^, Microplate Array Diagonal-Gel Electrophoresis (MADGE)^23^ and pulsed field electrophoresis^24^. Commercially, some all-in-one gel systems try to simplify the process like E-gel^25^, and some new dyes have reduced toxicity or increased brightness like GelRed.^26^ However, running agarose slab gels to separate nucleic acids is almost the same now as it was in the 1970s. Use of gel electrophoresis outside of a lab environment is rare due to general need for high voltages, large buffer volumes, and gel preparation. One notable exception is the E-gel system, which uses cartridge based “buffer less” gels and a power bank and reader to perform and analyze electrophoresis. We took some inspiration from this system to reimagine gel electrophoresis for our very specific application of DNA nanoswitch biosensing.

Most gel electrophoresis systems tend to be “one size fits all”, which can be useful for a lab environment but can also be wasteful and inefficient for targeted applications such as our DNA nanoswitches. Some other common uses of electrophoresis also only use a small subset of gel area, including validating PCR products^27,28^, plasmid insertion^29^, and DNA origami^30^. For these types of applications, the typical large format gel has many drawbacks including higher cost of equipment and reagents, higher running voltage and power consumption, and longer preparation times. Most of these drawbacks are driven by a single design choice – distance between electrodes. Closer electrode spacing reduces overall footprint/cost, volume, running voltages, and overall power.

At the heart of our innovations here is the enabling technology of 3D printing for prototyping and manufacturing. 3D printing enables affordable, rapid, on-site mass production in virtually any setting, and has become increasingly common in research^31– 33^. We have previously used 3D printing in our lab for development of the Centrifuge Force Microscope^34,35^, and others have used it to build light microscope^36^, centrifuge^37^, microfluidics^38^, rehabilitation device^39^ and other small laboratory equipments^40^. The printable electrodes allowed us to rapidly and inexpensively print and test various designs and electrode spacings.

Aside from performance considerations, the mini gel system has a large cost advantage over commercial systems. Lab grade gel electrophoresis equipment typically costs several hundreds of dollars for the gel box and can be tens of thousands if including a high voltage power supply and imaging equipment. These costs and electrical requirements of conventional gel electrophoresis equipment have prevented agarose gels from being realized as an option for use outside the lab environment.

This work narrows the gap to achieving point of care (POC) utility with DNA nanoswitches and a gel based readout. Several applications that have already been developed such as viral RNA detection may benefit from such a format, potentially enabling healthcare providers to conduct diagnostic tests and monitor patients’ health status in real-time, at or near the point of care. This would be particularly useful in remote and resource-limited settings where access to conventional laboratory facilities is limited. Our 3D printed custom gel box was produced for ∼$0.30 (and could likely be optimized to reduce cost below $0.10) and the assay can be run on battery power outside of a lab environment. Our lab system costs >$1,000 for reference with standard mini gel system and power supply each costing >$500. The 3D printed gel boxes can be used as disposable or can be reused multiple times, and it may be possible for gels to be pre-cast and vacuum sealed for immediate use at the point of care. Bands can be seen with as little as a blue flashlight and a cell phone camera. Further improvements in bulk manufacture, storage, transport, and user guidelines would still be needed, but we believe we have demonstrated that a gel readout can be a viable option.

Beyond point of care, we hope that our developments here will encourage other researchers to think “outside of the box” when considering electrophoresis. Just because it has been a one-size-fits-all lab technique that’s tried and true doesn’t mean it can’t be improved or made more efficient. There is still substantial room for innovation, and this work just scratches the surface. Customization of gel box size and shape, comb shapes and number of wells, and electrode position and orientation are all possible. These customizations may provide additional utility for various applications, without substantial additional cost.

With a gel box that can be easily and inexpensively reconfigured the to suit individual researchers’ needs, we see applications that certainly could go far beyond our own DNA nanoswitch sensor. We hope researchers might to consider, the next time they prepare liters of buffer and run gels in equipment costing thousands of dollars, whether such setups are truly ideal for their specific application or simply a matter of convention. This work demonstrates that thoughtful engineering and purposeful simplification can extend gel electrophoresis beyond traditional laboratory constraints and open new possibilities for accessible, application-driven use.

## Acknowledgements

The authors thank Jibin Abraham Punnoose and Arun Richard Chandrasekaran for useful discussions on the project. Research reported in this publication was supported by the National Institutes of Health through the National Institute of General Medical Sciences under award R35GM124720 to KH.

## Experimental Section

### Minigel Design and Fabrication

All Minigel models were designed using Autodesk Fusion 360 (version 2605.1.39). Models were exported in .stl format into Ultimaker Cura (v. 5.9.1) 3D printing software for formatting. Gcode files were exported and printed on a dual extruder Ultimaker 3 FFF 3D printer (PN:9671) equipped with two Ultimaker AA 0.4mm Print cores (PN:9529). All Minigel box models, unless otherwise specified, were prepared using default normal resolution (0.15mm) print settings for 2.85 mm Ultimaker White or Black Tough PLA (Poly Lactic Acid) with 100% infill, lines infill settings. Minigel electrodes were prepared using 2.85 mm ProtoPasta Electrically Conductive Composite PLA with these settings: nozzle temperature 230°C, 100% infill, lines infill settings. All Minigel systems were used without further post processing.

Portable power was supplied using alkaline batteries, including four Procell AAA 1.5 V cells, four Energizer LR44/A76 1.5 V button cells, and a Procell 9 V alkaline battery. These configurations were used to provide nominal output voltages of 6 V and 9 V for device operation. The battery power source was directly connected to the anode and cathode terminal posts of the Minigel using a pair of alligator clips with jumper wires.

### DNA Nanoswitch Reactions

All optimization experiments were performed using a previously reported DNA nanoswitch designed to detect Let-7b miRNA. The nanoswitch reactions, unless otherwise specified, were performed at 30ºC in 20 mM tris-HCl (pH 8) supplemented with 30 mM magnesium chloride (MgCl_2_) with equimolar amounts of Let-7b Nanoswitch and analyte (DNA mimic of Let-7b miRNA, 5’-TGAGGTAGTAGGTTGTGTGGTT-3’) with a reaction concentration of ∼400pM. Nanoswitch reaction mixtures were incubated for one hour, then 2:1 6X loading dye: 10X DNA stain (Biotium GelRed) was added to a final concentration of 1X and mixed. Ten microliters of sample was loaded to each well and electrophoresed at various voltage-times as specified in the results section. Total RNA was extracted from HeLa cells (BioChain, Cat# R1255811-50) and diluted to a final concentration of 250□ng/µL prior to use as input material for target detection with the DNA nanoswitch. All the buffers and reagents were prepared using nuclease-free water (Cytiva, Cat# SH30538).

### Agarose Gel Electrophoresis

All agarose gels used in all the experiments were prepared from Sigma-Aldrich agarose, Bioreagent low EEO (Cat. No. A9539) and freshly prepared 0.2 µm filter sterilized 0.5X tris borate EDTA buffer (TBE), unless otherwise specified, in 0.8% w/v ratio. 1kb DNA ladder from New England Biolabs (Cat# N3232) was used as reference sample. Standard agarose gels contained 25 mL of agarose gel were prepared and run on a Thermo Scientific Owl EasyCast B1A Mini Gel electrophoresis system (Cat# B1A-BP). Minigels were cast in 3mm thick slabs using variable volumes of agarose gel depending on the following volumetric ratio: 750 µL molten agarose per 10 mm gel length (Supporting Information, Table S1). All AGE, except battery powered Minigels, were run in 0.5X TBE running buffer at constant voltage using a Labnet International Inc. Enduro 300V power supply (Model: E0303) with the voltage-times specified in the results section.

### Gel Imaging

All gels images for optimization experiments were captured by an Azure Biosystems Azure 400 Bioanalytical Imaging System. Gels were imaged with ethidium bromide or SYBR Safe settings. Various exposures (500ms - 20s) were taken to maximize sensitivity while minimizing pixel saturation. Raw TIFF files were exported to Bio-Rad Image Lab software (V 6.1) for analysis. To capture the gel images by cell phone, the gel stained with SYBR Safe was illuminated by blue LED (Thorlab LED465E), and an orange safety filter was used.

