## Supplemental Information for "A 3D printed mini-gel electrophoresis system for rapid and inexpensive DNA nanoswitch biosensing"

^2^current address: School of Advanced Manufacturing and Robotics, Peking University, Beijing 100871, China

**Table of Contents:**

Figure S1. Effect of electrode-gel spacing on gel performance

Figure S2: Repeatability of mini gels

Figure S3: Use of acrylic vs 3D printed gel combs

Figure S4: Use of various gel percentages in resolving DNA nanoswitches

Table S1: Agarose gel and running buffer volume used in different sized box

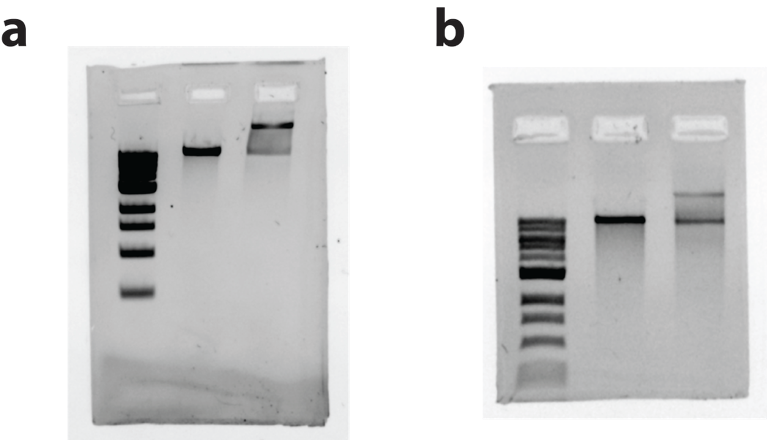

**Figure S1. Effect of electrode-gel spacing on gel performance.** a) Agarose gel run in a 40 mm minigel box with no gap between the electrodes and the gel. b) Agarose gel run with a 3 mm gap between the electrodes and the gel on each side. While both configurations enable electrophoretic separation, some visible differences are observed. The gel run without gaps exhibits slower migration and visible distortion of the gel matrix near the bottom edge, likely due to localized heating and partial melting. In contrast, the gel run with electrode-gel gaps shows slightly higher migration and no observable gel deformation.

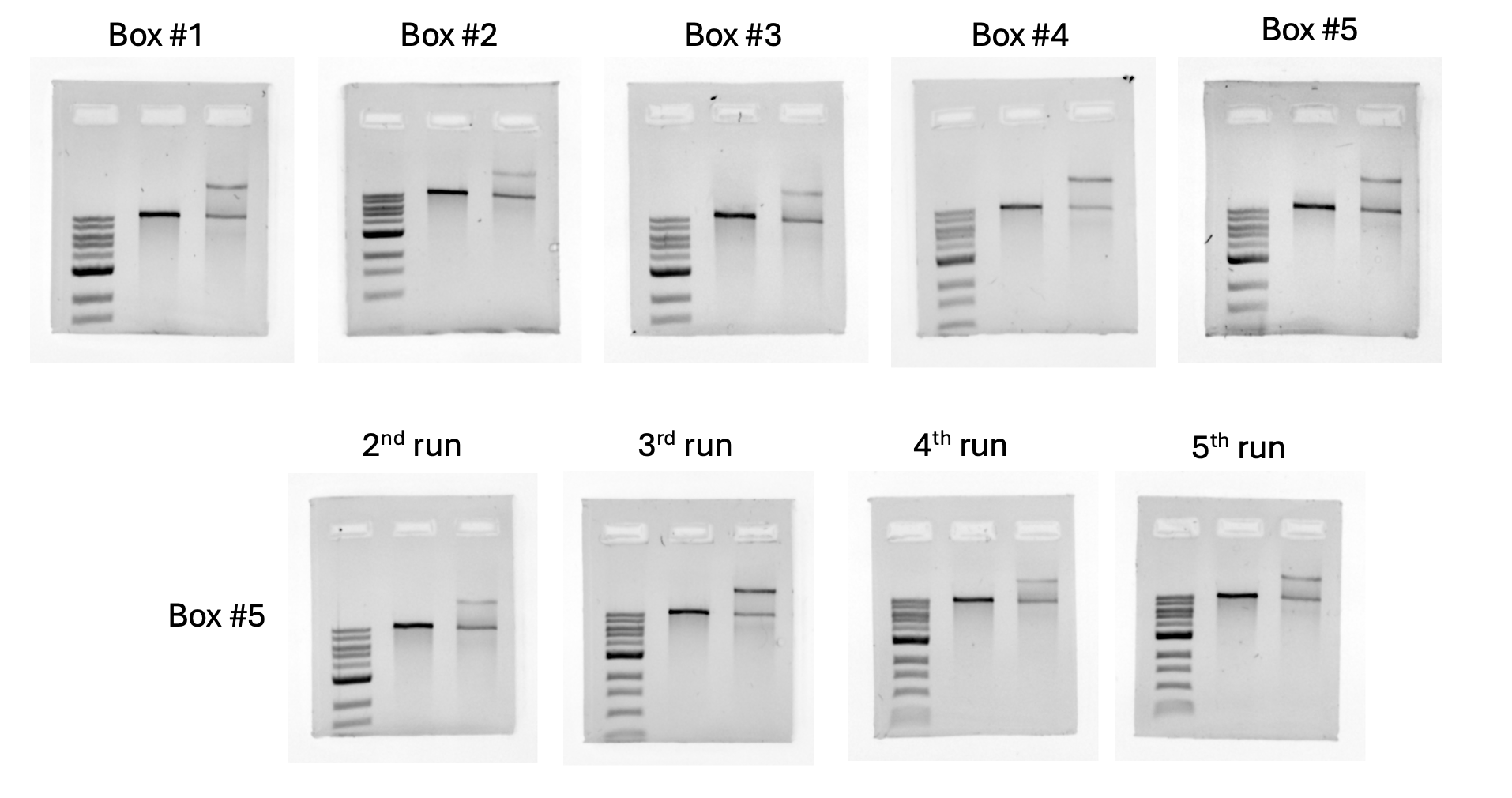

**Figure S2: Repeatability of mini gels.** Five separate gel boxes were printed and run for the first time, and box #5 was run for a total of 5 times.

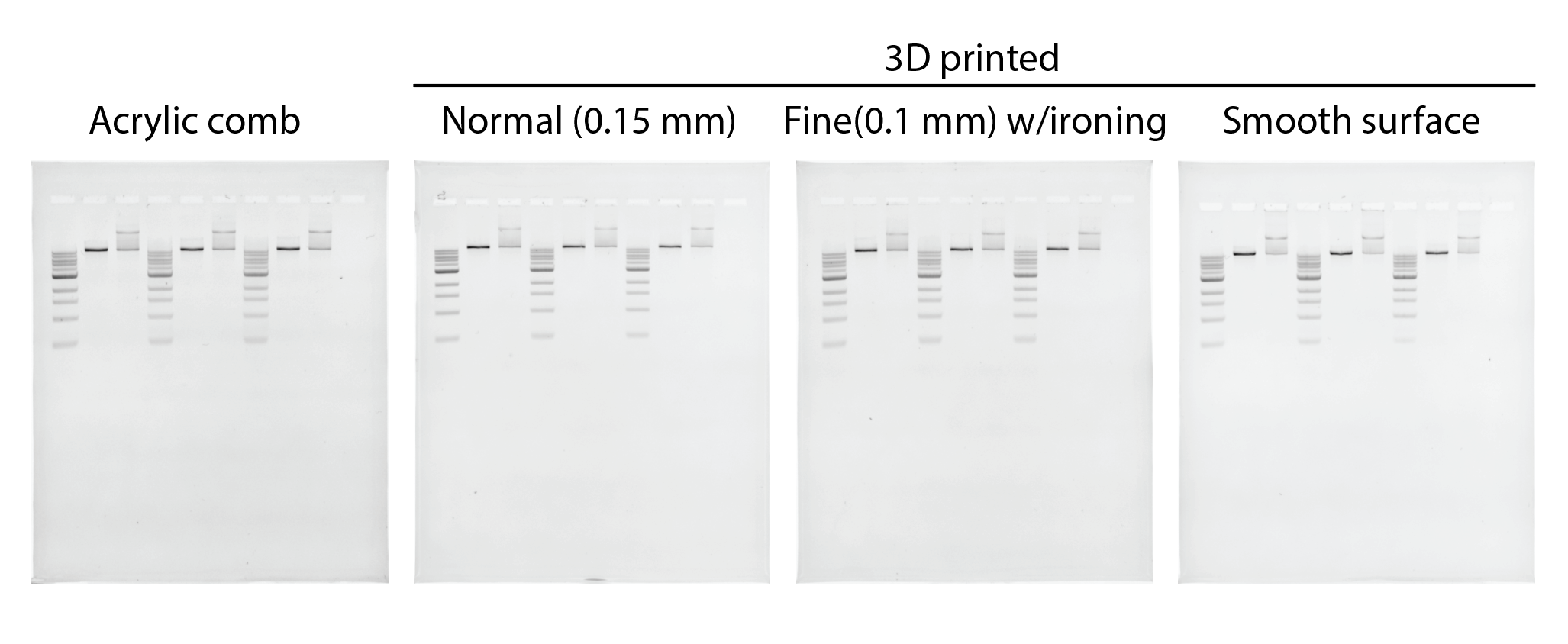

**Figure S3: Use of acrylic vs. 3D printed gel combs.**

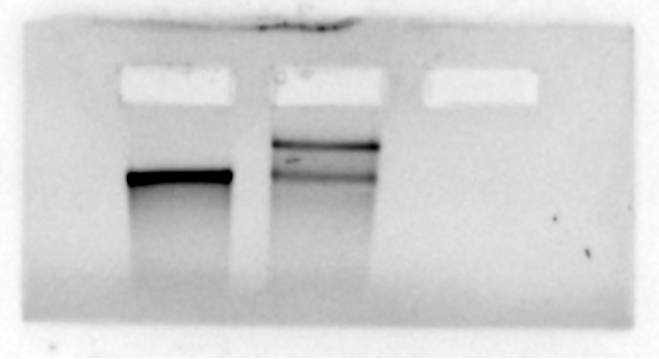

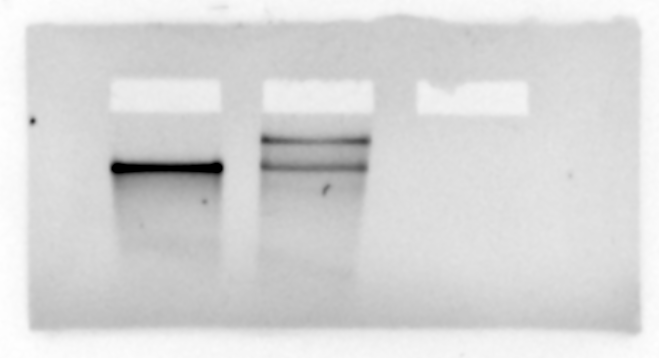

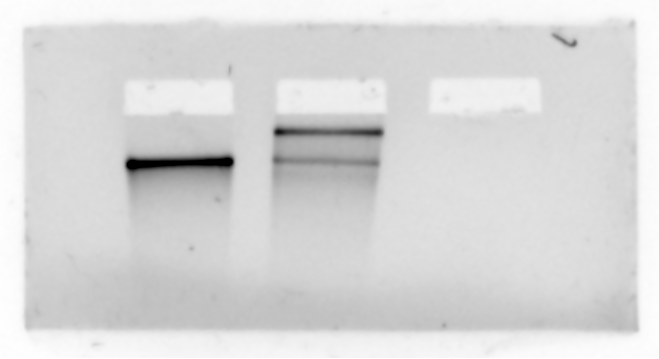

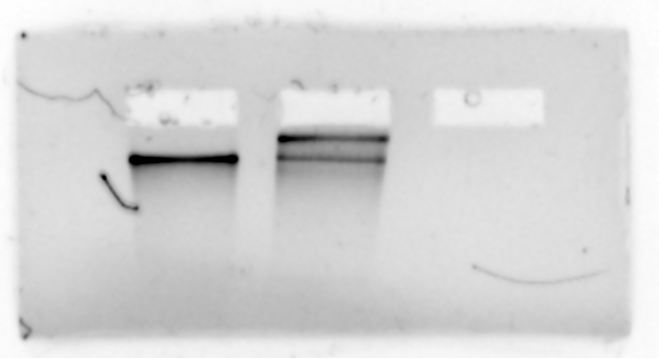

0.9 %

1.0 %

1.1%

1.2 %

0.7 %

0.8 %

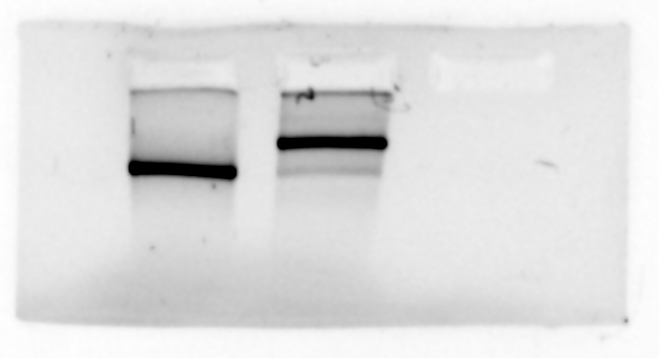

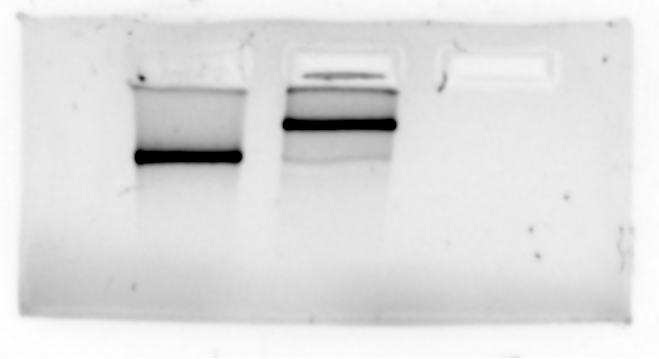

**Figure S4: Use of various gel percentages in resolving DNA nanoswitches.** Tested different gel concentration between 0.7% to 1.2% w/v with constant running conditions in 15mm gel box.

**Table S1. Agarose gel and running buffer volume used in different sized box**

| Sr. # | Box length (Inter electrode distance) | Agarose gel volume  (mL) | Running buffer used  (mL) |
| --- | --- | --- | --- |
| 1 | 137mm | 25 | 400 |
| 2 | 40mm | 3 | 3 |
| 3 | 35mm | 2.7 | 2.7 |
| 4 | 30mm | 2.2 | 2.2 |
| 5 | 25mm | 1.8 | 1.8 |
| 6 | 20mm | 1.5 | 1.5 |
| 7 | 15mm | 1.1 | 1.1 |
| 8 | 10mm | 0.75 | 0.75 |
